# High nucleotide diversity accompanies differential DNA methylation in naturally diverging populations

**DOI:** 10.1101/2022.10.08.511291

**Authors:** James Ord, Toni I. Gossmann, Irene Adrian-Kalchhauser

## Abstract

Epigenetic mechanisms such as DNA methylation (DNAme) are thought to comprise an invaluable adaptive toolkit in the early stages of local adaptation, especially when genetic diversity is constrained. However, the link between genetic diversity and DNAme has been scarcely examined in natural populations, despite its potential to shed light on the evolutionary forces acting on methylation state. Here, we analysed reduced-representation bisulfite sequencing and whole genome pool-seq data from marine and freshwater stickleback populations to examine the relationship between DNAme variation (between- and within-population), and nucleotide diversity in the context of freshwater adaptation. We find that sites that are differentially methylated between populations have higher underlying standing genetic variation, with diversity higher among sites that gained methylation in freshwater than those that lost it. Strikingly, while nucleotide diversity is generally lower in the freshwater population as expected from a population bottleneck, this is not the case for sites which lost methylation which instead have elevated nucleotide diversity in freshwater compared to marine. Subsequently, we show that nucleotide diversity is higher among sites with ancestrally variable methylation and also positively correlates with the sensitivity to environmentally induced methylation change. Both suggest that as selection on the control of methylation state becomes relaxed, so too does selection against mutations at the sites themselves. Increased epigenetic variance in a population is therefore likely to precede genetic diversification.

## Introduction

DNA methylation (DNAme) is an epigenetic mark whose roles in genome regulation, including gene expression regulation and transposable element suppression, have been well studied (He et al. 2011). Its role in local adaptation and long term evolution however, for example via plasticity, remains a topic of active debate. Methylome data support a potential role for DNAme in local adaptation in several species (Dubin et al. 2015; Sammarco et al. 2022), revealing that some genomic regions show differential methylation (DM) between different locally adapted populations. Populations (Johnson & Kelly, 2020) and species (Vernaz et al. 2021) with low genetic divergence from one another have been found to differ considerably in DNAme patterns at environmentally relevant loci, raising the possibility that flexibility of DNAme can be a source of phenotypic variation which increases adaptive potential of populations when genetic diversity is challenged. However, DNAme is far from independent of genetic diversity; it is determined by the presence of sites with the capacity to be methylated (typically a in a CpG context) and is subject to *trans*- and *cis*-regulation (Villicaña and Bell 2021). Therefore, the potential for methylation and corresponding plasticity is determined by the local genomic context, and thus the ‘epigenetic potential’ of a population evolves at the sequence level (Kilvitis et al. 2017). It is also established that the epigenetic conformation of the genome affects the propensity for sequence change. Notably, DNAme is well known to influence mutation rates due to the higher susceptibility of methylated Cs to spontaneous deamination to form thymine (Xia et al. 2012; Poulos et al. 2017; Zhou et al. 2020). Therefore, DNAme and sequence variation are intertwined (see also Adrian-Kalchhauser et al. 2020).

Despite the interdependence of DNAme and sequence variation, the potential importance of this link in local adaptation has been largely overlooked. For example, typical workflows for detection of differential methylation tend to exclude CpG sites that are not detected at a certain coverage in a certain proportion of individuals (e.g. Akalin et al., 2012), and therefore it could be assumed that genetic diversity of those sites is irrelevant. However, these sites may nevertheless harbour genetic variants in the population, the relative frequencies of which may be informative about evolutionary forces acting on methylation state, potentially allowing further dissection of the manner in which DM evolves in the context of local adaptation. Indeed, methylation sites within certain promoters have already been shown to exhibit selective sweep signatures in *Arabidopsis* (Shirai et al. 2021). Epigenetic diversification is one of many possible routes to local adaptation (e.g. Smith et al., 2016) but may occur in conjunction with others, such as selection on discrete new mutations (hard sweeps) or on standing genetic variation (SGV) (soft sweeps) (Bernatchez 2016; Hermisson and Pennings 2017). For example, if epigenetic modifications at multiple loci could confer similar adaptive benefit, epigenetic diversification could occur in conjunction with a soft sweep. Furthermore, given that methylated cytosines incur a heightened mutation risk, a gain in methylation in a divergent population may increase mutation rates at affected sites and *vice versa*. There is therefore a need to examine the relationships between differential methylation and nucleotide diversity in divergent populations.

The threespine stickleback fish (*Gasterosteus aculeatus*) has long been a popular model to study the genetics and, more recently, the epigenetics, of local adaptation. Ancestrally a marine fish, *G. aculeatus* has repeatedly and rapidly colonised freshwater habitats over the last few millennia, with large waves of colonisations having occurred since the formation of glacial lakes following the last ice age (Jones et al. 2012; Roberts Kingman et al. 2021). Freshwater-adapted morphs show numerous phenotypic adaptations including the loss of armour plating (Barrett et al. 2008) and changes to kidney morphology (Hasan et al. 2017). Phenotypic and genetic divergence has been observed over short time scales (Lescak et al. 2015), with large shifts in frequencies of particular alleles having been observed over just a few years in newly established lake populations (Roberts Kingman et al., 2021). Such rapid fixation of alleles on the basis of SGV is characteristic of a soft sweep (Bernatchez 2016). However, the rapid adaptability and high plasticity of sticklebacks (Day et al. 1994) also makes the contribution of epigenetic variation to freshwater adaptation compelling. Multiple studies have used bisulfite sequencing to reveal differentially methylated sites or regions between marine and freshwater populations, some of which are in the vicinity of genes relevant to freshwater adaptation (Smith et al. 2015; Artemov et al. 2017; Heckwolf et al. 2020; Hu et al. 2021).

A potential role for epigenetic variation in freshwater adaptation is especially pertinent given that the formation of freshwater populations has been characterised by population bottlenecks, constraining genetic diversity. Steep declines in the effective population size *Ne* have been observed in newly established freshwater populations both from time series experiments (Aguirre et al. 2022) and ancient DNA (Kirch et al. 2021). Interestingly, in their comparison of gill DNA methylomes between marine and freshwater fish in the White Sea region, Artemov et al. (2017) showed that freshwater fish had higher variance of DNAme compared to marine fish, in line with the idea that higher epigenetic variation could compensate for reduced genetic diversity, enhancing the adaptive potential of a population following a bottleneck.

While genome and / or epigenome data have been generated from multiple stickleback populations across the Northern hemisphere, the White Sea population complex is unique in that a variety of different data types have been generated from populations inhabiting the same region, including DNAme (Artemov et al., 2017), mRNA and small RNAseq (Rastorguev et al. 2016; Rastorguev et al. 2017), and whole genome pool-seq (Terekhanova et al., 2014; 2019).

Freshwater colonisation in this region has occurred relatively recently, with the oldest sampled lake estimated to have been formed approx. 700 years ago. Nucleotide diversity is likely to have decreased overall in this freshwater population due a past bottleneck (Terekhanova et al. 2014), but could be reduced further at specific sites due to purifying selection if the function of those sites becomes more important. However, diversity may increase due to lifting of selective constraints at sites whose function is less important in the new environment, or through locally elevated mutation rate due to methylation gain. Shifts in nucleotide diversity at DMCs may therefore be informative about the evolutionary forces acting on DNAme during local adaptation.

To address these intriguing hypotheses we combine epigenetic and genetic data from the White Sea stickleback population complex to study the interactions between methylation differences and nucleotide diversity during freshwater colonisation. We examined nucleotide diversity in relation to methylation divergence, variance, and the environmental inducibility of methylation state, considering both variance and inducibility as indicators of the relative stringency DNAme regulation.

## Results

### Elevated nucleotide diversity accompanies differential methylation but level depends on the direction of methylation change

For generation of DNAme and genome sequence data, respectively, both Artemov et al. and Terekhanova et al. sampled freshwater fish from the same lake (Lake Mashinnoye) and marine fish from nearby coastal locations in the Kandalaksha gulf. A combined analysis of samples from these two datasets therefore allowed us to identify differentially methylated cytosines in CpG context (DMCs) between marine and freshwater populations and examine the nucleotide diversity of these sites in separate samples of those populations (**Fig. 1**).

**Figure 1.**
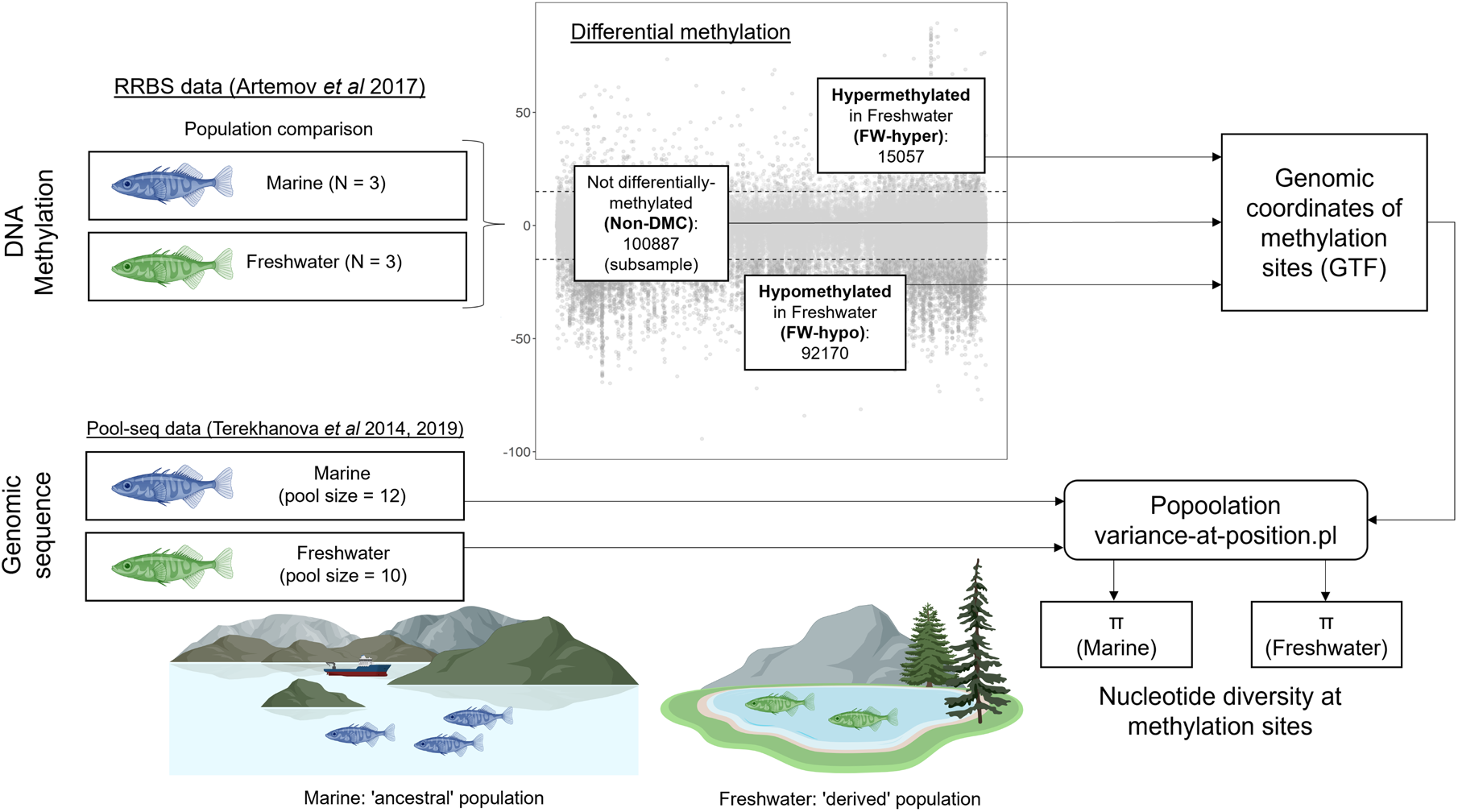
Analysis workflow for obtaining nucleotide diversity estimates for differentially methylated sites. Site-level differential methylation analysis was performed to compare marine (considered as ‘ancestral’ population) vs freshwater sticklebacks (considered as ‘derived’ population) using RRBS data previously published by Artemov *et al* (2017). Taking the marine population as the reference methylation state, sites were classified as FW-hyper (significantly higher % of methylated copies in freshwater compared to marine), not differentially-methylated (Non-DMC; no significant difference in % methylated copies between populations), or FW-hypo (significantly lower % of methylated copies in freshwater compared to marine). For non-DMCs, a subset of the total was used, comprising ∼11% of the total non-DMCs (see methods). Coordinates of sites belonging to the three site classes (Non-DMC, FW-Hypo, and FW-hyper) were compiled in a GTF file for use with the variance-at-position.pl script from the Popoolation toolkit. The nucleotide diversity (π) of each site class on each chromosome was estimated from whole-genome pool-seq data (Terekhanova *et al*, 2014 & 2019) of marine and freshwater fish derived from the same or similar geographic locations as those taken for the RRBS data.

We first derived measures of nucleotide diversity of differentially methylated (DMCs) and non-differentially methylated DNAme sites (Non-DMCs) in the form of π (average number of pairwise differences within population) (Nei and Li 1979), Watterson’s *θ* (Watterson, 1975), Tajima’s *D* (Tajima 1989), and pairwise Fst (Weir and Cockerham 1984). All nucleotide diversity measures were derived from pool-seq data at methylation sites detected from RRBS data (**Fig. 1**). In a site-level differential methylation analysis, we used a standard workflow to detect DMCs in RRBS individuals which retained sites for which a methylation call was obtained in all of the analysed individuals. These included six individuals as part of the main population comparison and a further five which were used for a subsequent analysis of site inducibility. Therefore, our analysis relied on the assumption that even sites that are conserved across RRBS individuals would show some level of genetic diversity in the slightly larger subsets of the populations profiled by pool-seq. Considering Non-DMCs as representing a baseline level of nucleotide diversity at cytosines in CpG context, we compared the π of Non-DMCs with that of DMCs that either lost methylation (significantly fewer methylated copies, FW-hypo) or gained methylation in freshwater compared to marine (significantly more methylated copies, FW-hyper). The Non-DMCs we considered here represented a subsample of the total number of Non-DMCs which was extracted by taking one random Non-DMC within 2Mb of each DMC.

We observed consistent associations between differential methylation and nucleotide diversity. DMCs showed elevated π compared to Non-DMCs regardless of population, with the highest π observed amongst sites which gained methylation in freshwater (FW-hyper) (**Fig. 2A**). When comparing π between populations for each category of sites, π was slightly though significantly reduced in freshwater compared to marine at Non-DMCs, reflecting the expected reduction in nucleotide diversity in the derived population. π calculated for sliding windows across chromosome 1 suggested that the pattern of reduced diversity in freshwater is a genome-wide feature (**Fig. S1A**) which likely resulted from a past bottleneck. Freshwater π was similarly lower at FW-hyper sites, but not at FW-hypo sites which rather showed elevated π in freshwater compared to marine. Watterson’s *θ* showed a similar pattern to that of π (**Fig. S2**). Patterns of between-population diversity (Fst) were not so pronounced: When compared to Non-DMCs, FW-hypo sites had slightly increased Fst, but FW-hyper sites did not (**Fig. 2B**). Tajima’s *D* was mostly negative in both populations as indicated by sliding windows across chromosome 1 (**Fig. S1B**), but was relatively higher in freshwater, consistent with the scenario of a population bottleneck following freshwater colonisation. Tajima’s *D* of methylation sites largely reflected this chromosome-wide pattern of higher values in freshwater, with the exception of FW-hyper sites which did not differ in Tajima’s *D* between the two populations (**Fig. 2C**). Within populations, Tajima’s *D* tended to be higher among DMCs than non-DMCs, with the exception that in freshwater, FW-hyper sites showed no difference in Tajima’s *D* compared to Non-DMCs.

**Figure 2.**
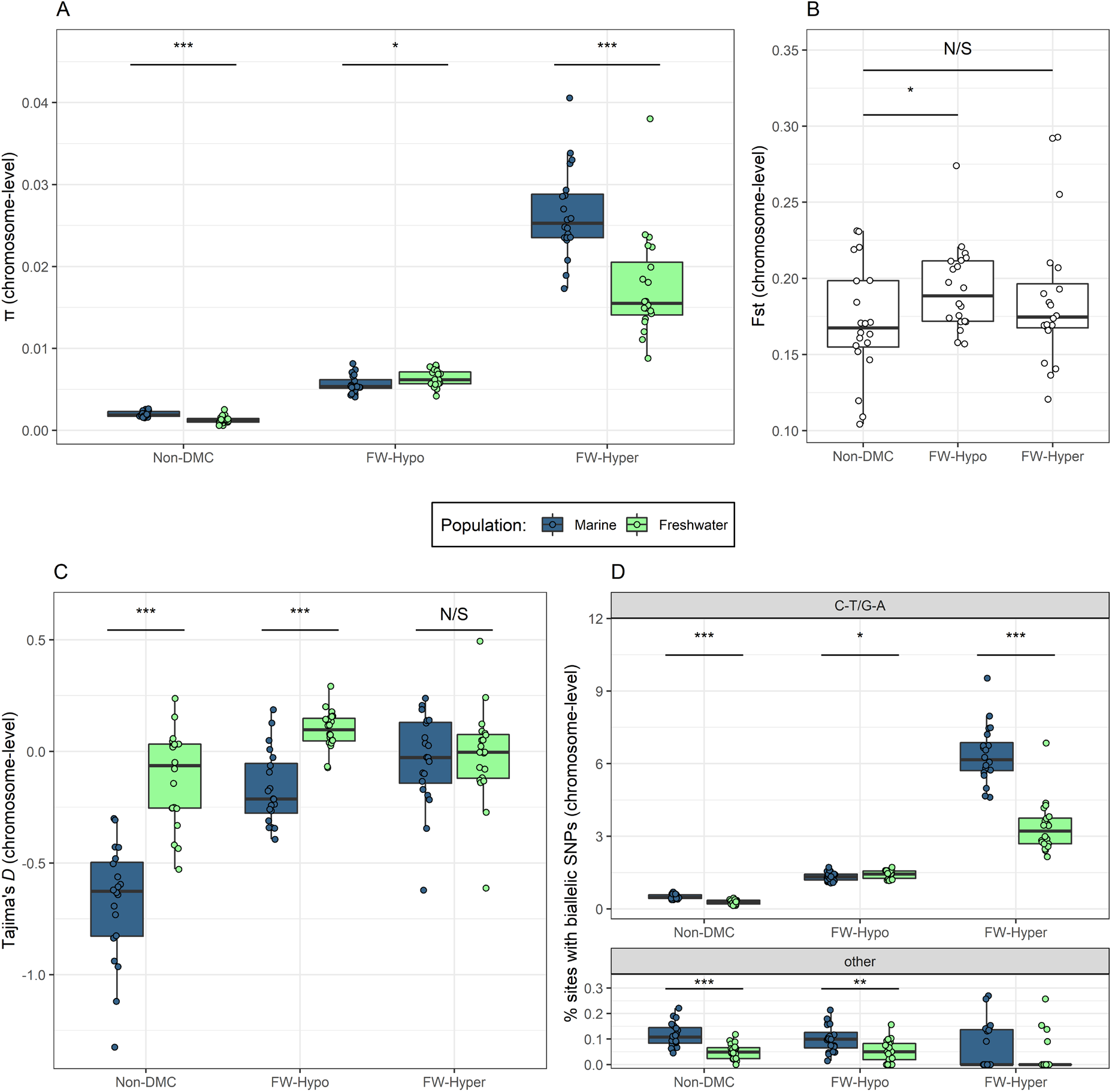
Nucleotide diversity of differentially methylated sites. **(A)** π (average number of pairwise differences), **(B)** Fst (marine vs. freshwater), and **(C)** Tajima’s *D* estimated from pool-seq of marine and freshwater sticklebacks for three classes of methylation site identified from RRBS individuals and classified according to the direction of methylation difference in freshwater fish compared to marine (Not differentially-methylated (Non-DMC), hypomethylated in freshwater (FW-Hypo), or hypermethylated in freshwater (FW-Hyper)). Each measure was estimated for each chromosome separately, such that one point represents an estimate from one chromosome. Significance stars derive from paired Wilcoxon tests (comparison of chromosome pairs). **(D)** % of sites in each site class harbouring biallelic SNPs of the type C-T/G-A (upper panel) or other types (C-A/G-T or C-G/G-T, lower panel), estimated separately for each chromosome. Significance stars derive from paired *T*-tests.

These patterns of elevated nucleotide diversity were not driven by enrichment for DMCs in regions of high diversity. Rather, elevated π of DMCs was found to be strongly localised around individual DMCs (**Fig. S3**). The pattern was largely consistent across different genomic features including CpG islands, gene bodies, promoters, and intergenic regions, with the exception of DMCs which fell within DMRs (**Fig. S3**).

Next, we determined which mutation type(s) were most likely to be driving the elevated π of DMCs, and specifically whether this was driven by an over-abundance of C-T transitions. To this end, the percentages of sites in each category harbouring biallelic SNPs of different types (C-T/G-A, C-A/G-T, or C-G/G-C) were calculated from the pool-seq data. The majority of SNPs were C-T/G-A, comprising 92% of SNPs across all categories in marine and 95% in freshwater. The proportion of sites harbouring biallelic C-T/G-A SNPs across the three categories of methylation site and two populations showed essentially an identical pattern to that of π, with FW-hyper sites harbouring the highest proportion of C-T/G-A SNPs in both populations (**Fig. 2D**). Marine had more C-T/G-A SNPs than freshwater in the Non-DMC and FW-hyper categories but not the FW-hypo category, in which the freshwater population had more C-T/G-A SNPs, also consistent with the pattern observed in π. Meanwhile, the % of other SNP types showed no clear differences between the site categories and was not elevated in the freshwater population amongst FW-hypo sites. Therefore, both the general increased nucleotide diversity amongst DMCs and the heightened diversity of FW-hypo sites in freshwater seemed to be driven by a greater occurrence of C-T mutations.

### Higher diversity of infrequently-methylated DMCs

The finding that sites which gained methylation in freshwater (FW-hyper) had the highest π and highest proportion of C-T/G-A mutations population (**Fig. 2**) was contrary to expectations, as these sites would be expected to be infrequently methylated in the ancestral population and therefore not at high risk of mutation via deamination. We therefore tested the relationship between π and the distributions of % mean methylation across the three site categories. Here, mean % methylation refers to the average % of copies on which the C is methylated, or in other words the average frequency of the methylation mark among copies of given CpG site.

Because pool-seq data are not appropriate for estimating π at the level of a single site, we used a ranking procedure to examine the relationship between methylation frequency and π. Sites were divided into ranks according to their mean per-site % methylation, with higher ranks containing sites with higher % methylation. This ranking was performed separately for each population and each site category, and a measure of π obtained for each rank. We found that Non-DMCs displayed a bimodal density distribution, with most sites either very frequently or very infrequently methylated (**Fig. 3A**, left). The relationship between rank mean % methylation and π was clearly non-monotonic for non-DMCs, with the highest values appearing at low to intermediate methylation of around 25% (**Fig. 3B**, left). Meanwhile, the distributions of % methylation of DMCs were markedly different to those of Non-DMCs. FW-hypo sites were not characterised by a complete loss of methylation in the FW population. Instead, they were characterised by a shift from mostly high % methylation in marine to mostly intermediate % methylation in FW (**Fig. 3A**, middle). The higher π of FW-hypo sites in Freshwater appeared to be driven largely by sites in the low-intermediate range (**Fig. 3B**, middle). Meanwhile, FW-hyper sites were not characterised by a gain of methylation at previously bare sites, but instead were characterised by a shift from low-intermediate % methylation in marine to high % methylation in FW (**Fig. 3A**, right). Amongst FW-hyper sites, those with the highest methylation clearly contributed to the lower π of these sites in the freshwater population (**Fig. 3B**, right). In addition to absolute rank in % methylation, we also examined the relationship of π with the mean change in % methylation (i.e. the extent of hypo- or hypermethylation). We observed that among FW-hypo sites, the freshwater population had the largest increases in the π where the hypomethylation was strongest (**Fig. 3C** and **Fig. 3D**). FW-hyper sites also increased in π with the extent of hypermethylation (**Fig. 3C**), but this also corresponded with greater loss of π in FW (**Fig. 3D**). Meanwhile, sites with stronger methylation change in either direction (methylation loss or gain) had higher Fst (**Fig. 3E**).

**Figure 3.**
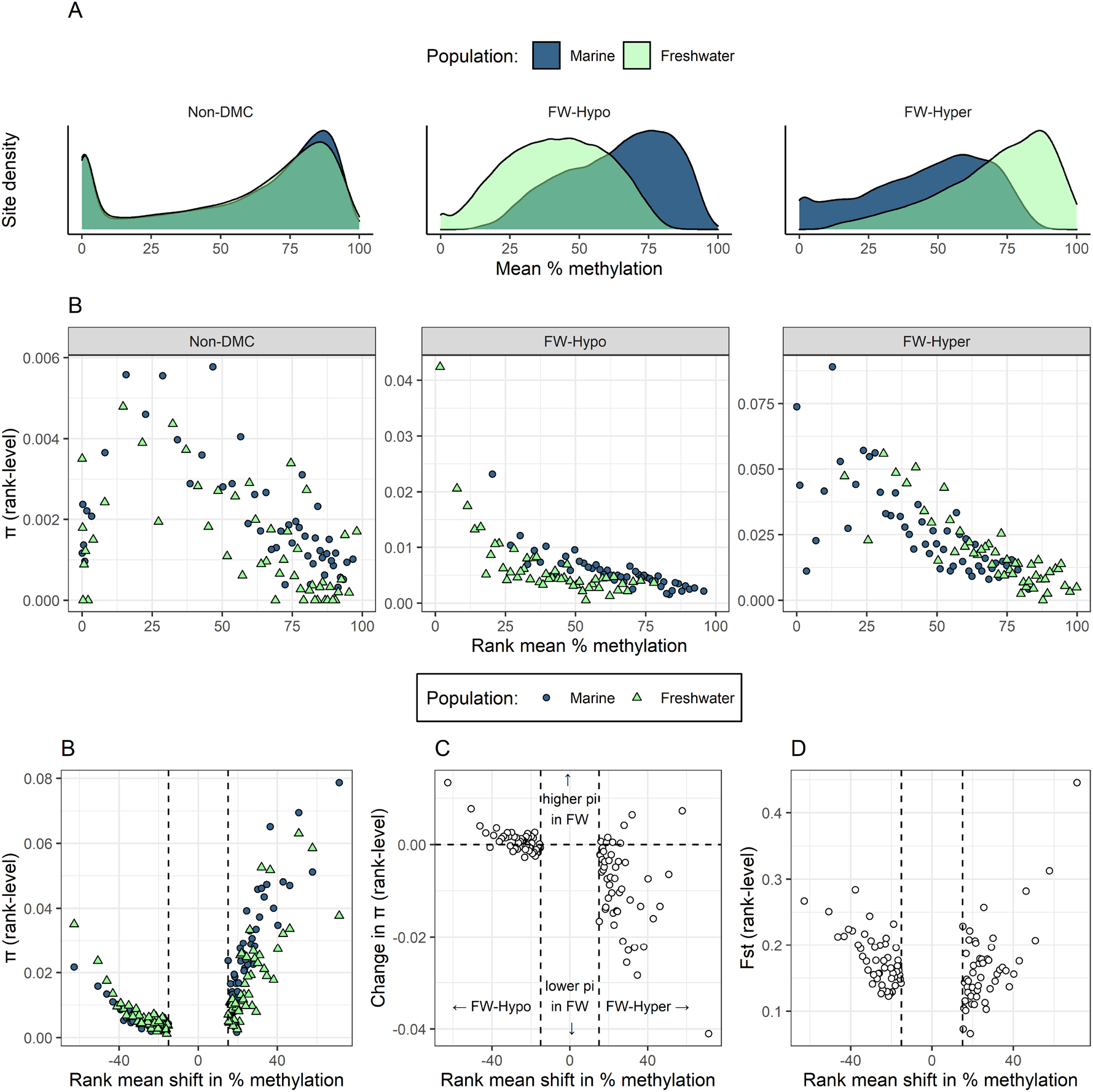
Distribution of mean % methylation (% of methylated copies at a given CpG site) in three site categories and their relationship with nucleotide diversity in marine and freshwater populations. **(A)** The relative distribution of sites along the mean % methylation in each population and site category. **(B)** π of sites that were ranked according to their mean % methylation in three site categories. A single π estimate was derived for all the sites in a given rank and π is plotted against the mean % methylation of sites in the rank. Sites were ranked separately for marine (blue circles) and freshwater (green triangles) populations and for Non-DMC (left panel), FW-hypo (middle panel), and FW-hyper sites (right panel). Each rank contains an average of 2038, 1843, and 301 sites for Non-DMC, FW-hypo, and FW-hyper, respectively. **(C-E)** Nucleotide diversity **(C)**, difference in nucleotide diversity (π of freshwater – π of marine) **(D)**, and pairwise FST (marine vs. freshwater) **(E)** of sites that were ranked according to the degree of hypo- or hypermethylation in freshwater. For each population, a single π estimate was derived for all the sites in a given rank. Each rank contains an average of 1843 and 301 sites for FW-hypo and FW-hyper sites, respectively.

### High nucleotide diversity accompanies high variability in ancestral methylation

Considering that sites with intermediate mean methylation frequency are liable to have more variable methylation levels than those with very low or very high methylation frequency, we also considered the relationship between π and the standard deviation (SD) of % methylation. We predicted that sites with more variable methylation would have higher nucleotide diversity, reasoning that the methylation state of these sites is not stringently controlled and therefore mutations at these sites may have little impact on function. We first examined the distributions of standard deviations of % methylation for sites in the Non-DMC, FW-hypo and FW-hyper categories. Non-DMCs were largely invariable, with slightly more variable methylation in freshwater compared to marine (**Fig. 4A**, left), consistent with the observations of Artemov *et al* (2017). FW-hypo sites were characterised by a pronounced increase in SD from ancestral to derived population, shifting from a left-skewed distribution in marine to a Gaussian-like distribution in freshwater (**Fig. 4A**, middle). Meanwhile, FW-hyper sites, showed a skewed distribution in marine with an elongated plateau to the right, while methylation was less variable in freshwater (**Fig. 4A**, right).

**Figure 4.**
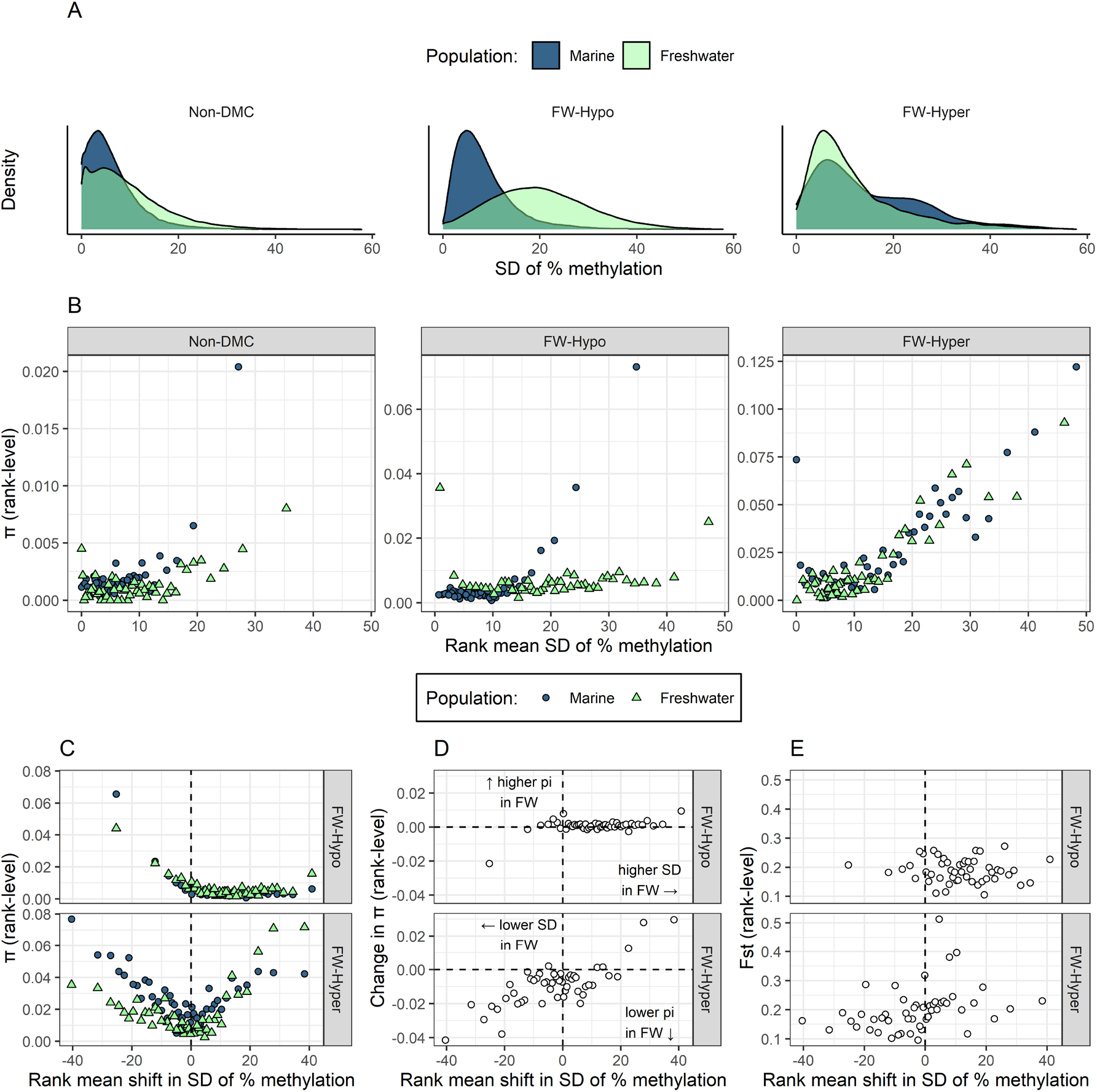
Distribution of standard deviation (SD) of % methylation levels in three site categories and two populations and their relationship with nucleotide diversity. **(A)** The relative distributions of SD of % methylation in each population and category. **(B)** π of sites that were ranked according to their SD of % methylation level. A single π estimate was derived for all the sites in a given rank and this rank-level π is plotted against the mean SD of sites in the rank. Sites were ranked separately for marine (blue circles) and freshwater (green triangles) populations and for Non-DMC (left panel), FW-hypo (middle panel), and FW-hyper sites (right panel). Each rank contains an average of 2038, 1843, and 301 sites for Non-DMC, FW-hypo, and FW-hyper sites, respectively. **(C-E)** π **(C)**, difference in π (π of freshwater – π of marine) **(D)**, and pairwise Fst (marine vs. freshwater) **(E)** of sites that were ranked according to difference in SD between marine and freshwater (SD of freshwater – SD of marine), where higher values represent higher SD in freshwater and lower values represent lower SD in freshwater. For each population, a single π estimate was derived for all the sites in a given rank. Each rank contains an average of 1843 and 301 sites for FW-hypo, and FW-hyper sites, respectively.

To examine the relationship between π and the variability in methylation, sites were ranked according to the SD of % methylation in each population and site category. We observed that π increased steeply at an SD above ∼15 in all three site categories in the marine population and two of the three site categories in freshwater (**Fig. 4B**). For the π of FW-hypo sites in FW however, there was no obvious relationship with the exception of two ranks showing highly elevated π at opposite ends of the range of SD values (**Fig. 4B**, middle). The high π of the lowest rank, which contradicted the trend observed in the other categories, is likely attributable to high ancestral diversity of sites that have almost completely lost methylation in freshwater (and therefore attain very low variance) but retain high ancestral nucleotide diversity. The relationship between π and the shift in SD of % methylation would appear to support this notion because sites with the largest decrease in SD in freshwater also have the highest π in marine (**Fig. 4C**). Shifts in the SD of methylation were also associated with shifts in π (**Fig. 4D**). Both amongst FW-hypo and FW-hyper sites, there was a trend towards reduced π with decreased SD and increased π with increased SD. No clear relationships were observed between the shift in mean % methylation and Fst for either FW-hypo or FW-hyper categories (**Fig. 4E**).

### Environmentally inducible DNA methylation is linked with higher nucleotide diversity

So far, we have considered differential methylation in regard to losses or gains of methylation that have been detected in a population ∼700 years after its colonisation of a new environment. While such changes may result from long-term processes, changes in methylation state can also be directly induced by the environment. Such inducibility may be important for adaptation but also subject to genetic variation. We therefore analysed additional samples from the dataset by Artemov et al. (2017), as in addition to the main population comparison, the authors also quantified gill methylation changes in fish from each population in response to a change in environmental salinity. Fish from each population were exposed to the opposite conditions, with marine fish exposed to reduced salinity and freshwater fish exposed to increased salinity (**Fig. 5A**). Here, we reasoned that environmental inducibility of methylation state could also reflect the degree of genetic versus environmental control of methylation state, similar to increased SD of methylation possibly reflecting relaxed control of methylation state. Sites whose methylation state is more responsive to the environment could be assumed to be under looser genetic control and possibly relaxed selection. In total, sites that were induced in either population constituted 3.9% of Non-DMCs, 12% of FW-hypo and 40% of FW-hyper sites, thus representing a substantial enrichment of induced sites among DMCs and FW-hyper sites especially. When considering the proportions of induced sites in each population separately (**Fig. 5B**), freshwater had a higher proportion of induced sites than marine among FW-hypo sites and a lower proportion among FW-hyper sites, a pattern that almost perfectly mirrored that which was observed for π (**Fig. 2A**) and C-T/G-A mutations (**Fig. 2C**).

**Fig. 5.**
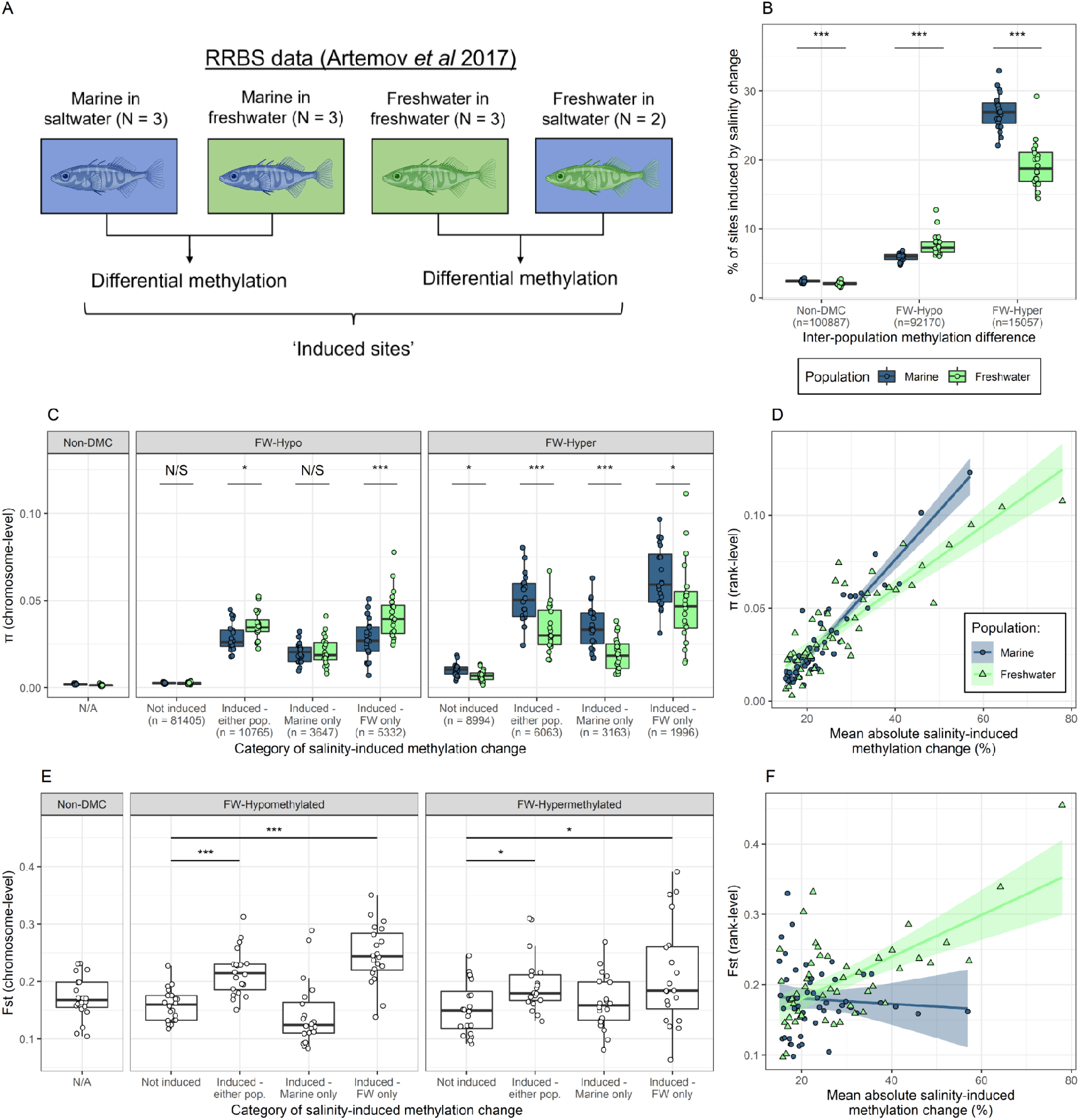
Nucleotide diversity of differentially methylated sites in relation to their capacity for induced methylation change in response to environmental salinity. **(A)** Additional RRBS data deriving from experimental salinity treatments performed by Artemov et al (2017) (marine fish placed in freshwater and freshwater fish placed in saltwater) were used to identify sites that were inducible in response to salinity change in either of the two populations. **(B)** % of sites in the Non-DMC, FW-hypo, and FW-hyper categories that were induced in response to salinity change in the marine (blue) and freshwater (green) populations. **(C)** Per-chromosome estimates of π for FW-hypo and FW-hyper sites divided according their capacity for induced gill methylation change in response to a change in environmental salinity, considering sites that were induced in neither of the populations, either of the two populations, or only in one of the two populations (marine or freshwater). π of Non-DMCs is shown in separate panel for comparison. **(D)** π of inducible DMCs that were ranked according to their mean absolute induced change in % methylation (i.e. regardless of the direction of the change). Within each population, only sites that were significantly differentially methylated in response to salinity change (mean change in % methylation >/= 15) were considered. Separate ranks were obtained for marine and freshwater populations and a single π estimate obtained for each rank. Each rank contains an average of 238 sites for Marine and 242 sites for Freshwater. **(E)** Per-chromosome estimates of pairwise Fst (freshwater vs. marine) of FW-hypo and FW-hyper sites divided according to their capacity for induced methylation change. Fst of Non-DMCs is shown in separate panel for comparison. **(F)** Pairwise Fst of inducible DMCs that were ranked according to their mean absolute induced change in % methylation (i.e. regardless of the direction of the change).

We find that sites that were induced in either population, or that were induced only in one of the two populations, had elevated π compared to sites that were not induced in either population (pairwise *T*-tests, all *P* < 0.01) (**Fig. 5C**), while the π of non-induced DMCs was closer to that of Non-DMCs. Furthermore, the gain in π in freshwater among FW-hypo sites appeared to be driven by sites that had gained inducibility (i.e. which were not previously inducible by environment), as this elevated π was observed among sites that were induced only in freshwater, but not sites that were induced only in marine. Meanwhile, amongst FW-hyper sites, π was consistently lower in freshwater compared to marine, regardless of the induced methylation category. When induced sites in each population were assigned to ranks according to the mean absolute induced methylation change (i.e. regardless of whether salinity change induced lower or higher methylation), π increased linearly with the mean induced methylation change in both marine (linear model, *R*^2^ = 0.87, *P* < 0.001) and FW populations (*R*^2^ = 0.76, *P* < 0.001) (**Fig. 5D**), suggesting that environmental inducibility can reliably predict nucleotide diversity and vice-versa. A linear model of π as a function of inducibility and population revealed a significant interaction between the two predictors (F(1) = 19.4, *P* < 0.001), reflecting the steeper relationship observed in the marine population.

Compared to non-induced sites, sites that were induced had significantly elevated pairwise Fst in both FW-hypo and FW-hyper categories (**Fig. 5E**). This elevated Fst was driven by sites that were induced in freshwater fish, as sites induced only in marine fish did not show elevated Fst compared to non-induced sites. The elevated Fst of FW-hypo sites (as seen in **Fig. 2B**) was therefore likely driven by sites that are inducible in the freshwater population. Finally, when induced sites in each population were ranked according to the degree of induced methylation change in that population, sites that were induced in the freshwater population showed a weak positive correlation between inducibility and Fst (linear model, *R*^2^ = 0.39, *P* < 0.001), while sites that were induced in the marine population did not show such a correlation (*R*^2^ = -0.02, *P* = 0.63) (**Fig. 5F**).

## Discussion

Here, we examined the relationship between DNA methylation changes and nucleotide diversity in an ancestral and a derived population of wild threespine stickleback. For this purpose, CpG sites with different methylation status in freshwater compared to the marine population were considered as changes in DNAme that occurred in the course of freshwater colonisation. Our analyses show that genetic diversity is intimately linked to variation in DNAme across both long (population differentiation) and short timescales (environmental responses). This link between DNAme and nucleotide diversity can shed light on the evolutionary forces acting on methylation state and could hint at the extent to which epigenetic changes precede sequence evolution.

### Sites prone to methylation divergence have high standing genetic variation driven by hypermutability of 5mC

Despite applying stringent filtering to retain only Cs in CpG context that were detected in all RRBS individuals (requiring all individuals to be either C/C or heterozygous at reference CpG loci), we found that not only did DMCs harbour SNPs among the individuals represented in the pooled sequencing dataset, but were even enriched for them. In other words, DNAme changes occurred at sites of high SGV. Consistent with a probable past bottleneck (Terekhanova et al. 2014), nucleotide diversity was reduced in freshwater, a pattern that held for FW-hyper sites despite the expectation that sites gaining methylation should incur higher mutation rates (Xia et al. 2012). However, FW-hypo sites – those that had lost methylation in freshwater – exhibited an increase in π in freshwater compared to marine, implying relaxed selection among these sites in the derived population.

The use of various measures of genetic diversity in parallel can provide insights into possible evolutionary processes at a fine scale. As well as higher π, DMCs also had higher Tajima’s *D* (higher proportion of intermediate-frequency alleles) than non-DMCs within the marine population, suggesting that sites with a tendency to diverge in methylation state are those that were already under weaker selective constraint (Jackson et al. 2015). The generally higher Tajima’s *D* in FW was expected following a recent population bottleneck (Stajich and Hahn 2005). However, this increase was not observed at FW-hyper sites, suggesting that while FW had an overall tendency to accumulate intermediate frequency alleles, this was impeded at FW-hyper sites, possibly due to increased selective constraint. Meanwhile, Fst was slightly elevated among FW-hypo sites compared to Non-DMCs, suggesting that sites that lost methylation are also subject to some degree of differential selection. Fst was not elevated at FW-hyper sites, however, despite evidence of their differential selection from the Tajima’s *D* statistic. This discrepancy may be explained by the high SGV of these sites, such that within-population diversity remains high despite a relative depletion of intermediate frequency alleles in the freshwater population. Contrasting the different measures of nucleotide diversity therefore reveals complex patterns of sequence evolution that depend upon both DNAme context and SGV.

DNAme and mutations rates are instrinsically linked by the hypermutability of 5mC. Indeed, we find that the patterns of nucleotide diversity are driven by higher abundance of C-T/G-A SNPs amongst DMCs, but not other SNP types, suggesting that they are driven by higher rates of spontaneous deamination of methylated Cs (Xia et al. 2012). The hypermutability of 5mC may further explain why FW-hypo sites were >6x more common than FW-hyper sites. Sites that acquire methylation during the transition to a new environment are more prone to mutation, and thus loss of the recently gained methylation, than sites losing methylation during the transition. Thus, stable gains in methylation are more difficult to attain than stable losses, and active selection might be required for their maintenance. Indeed, as sites with newly gained methylation must escape deamination in several individuals in order to be detected in the differential methylation analysis (due to stringent site filtering), FW-hyper sites may be enriched for the subset of new methylations that are under selective constraint.

### Relationships between nucleotide diversity and methylation frequencies further imply differential selection of DMCs

The relative frequencies at which sites are methylated (expressed as % methylation) capture both intra- and inter-individual variation in methylation state. Although the RRBS data derived from a specific tissue – gill, such a tissue is nevertheless heterogeneous, comprising of different specialised cell types (Pan et al. 2022). Very high or very low methylation frequencies are therefore likely to comprise sites where the same state is consistently maintained across the majority of cells and/or cell types. Although cell type-specific methylation is likely to be important in some contexts (Loyfer et al. 2022), it could nevertheless be inferred that sites with consistent methylation state are more likely to be selectively constrained. We observed that among Non-DMCs, most sites had either very high or very low % methylation, suggesting most sites have a methylation state that is consistently maintained across cell types (i.e. consistently methylated or non-methylated). Non-DMCs with intermediate methylation frequencies had higher π than those with very high or very low frequencies, again indicating stronger selective constraints on sites which are consistently either methylated or non-methylated. This is consistent with previous observations that sites in the human genome with low to intermediate methylation frequency *in vitro* have higher mutation rates in human populations (Xia et al. 2012). We found that in sticklebacks, differential methylation in the freshwater population was characterised by shifts either towards (FW-hypo) or away from (FW-hyper) intermediate methylation frequency. Accordingly, π increased in freshwater with the degree of hypomethylation and decreased with the degree of hypermethylation, while Fst tended to increase with the degree of change in either direction. Combined, these patterns imply that differential methylation occurs alongside differential selection, in that (stable) gains in methylation tend to be selectively constrained while sites that lose methylation are released from selective constraint. This would make sense given that a loss of methylation relinquishes the requirement of the locus to remain as a CpG dinucleotide.

### Genetic variation reflects (ancestral) epigenetic variation

Inter-individual variability of DNA methylation remains understudied in natural populations, and yet it may hint at the processes by which the methylome evolves. Using the same RRBS dataset, Artemov et al. (2017) previously showed that methylation was more variable in the derived freshwater population. Here, we show that this effect depends on differential methylation. Indeed, the higher variability in freshwater appeared to be driven largely by FW-hypo sites which showed a Gaussian-like distribution of SD values in the FW population compared to a strongly left-skewed distribution of the same sites in marine. FW-hyper sites on the other hand became less variable in FW. Loss of methylation in FW therefore appears to be characterised by relaxed control of methylation state, while gain of methylation is associated with tighter control. Indeed, heightened variability of FW-hypo sites suggests that loosening of regulation is itself the cause of methylation loss.

π tended to increase with variability of methylation, which would support the hypothesis that sites with less tightly maintained methylation state are under weaker selective constraint. This pattern was consistent across all three site categories in the marine population, but was absent among FW-hypo sites in FW. Genetic variation therefore reflects ancestral, but not recently acquired variability in methylation state. Relaxed selection on sites which lose methylation would lead to an accumulation of C-T mutations, while stronger selective constraint would reduce the nucleotide diversity of sites with stable methylation gain. Concordantly, sites that became less variable in freshwater tended to lose π, while sites that became more variable tended to gain pi. The pattern was more prominent among FW-hyper sites which predominantly had both decreased SD of methylation and decreased π. Among FW-hypo sites, the lack of correlation between SD and π in freshwater (or change in SD and change in π) could be explained by the relatively young age of the freshwater population (∼700 years) and subsequent lack of time for mutations to accumulate.

While it is plausible that the increased π and SD of FW-hypo sites reflects relaxed selection on the regulation of methylation state, an alternative hypothesis is that variable loss of methylation reflects epigenetic responses that have occurred only in a fraction of individuals in the population. This scenario would also be consistent with elevated π of the FW-hypo sites; if these differential epigenetic responses were genotype-specific, the elevated π would reflect SGV as opposed to new mutations. Such a scenario would be consistent with a soft sweep (Hermisson and Pennings 2017), in which selection could act on many different genetic and epigenetic loci, thus maintaining diversity at both levels.

### Environmental inducibility of DNA methylation may predict sequence evolution

The potential importance of epigenetic mechanisms in mediating plastic responses has long been discussed (Johnson & Tricker, 2010) and, more recently, demonstrated experimentally (Stajic et al. 2019). Although the environmental induction of a particular epigenetic state (e.g. addition or removal of DNAme) can occur in the context of adaptive mechanisms (Lämke and Bäurle 2017), such an induction does not always necessarily constitute an adaptive response. As such, we considered environmental inducibility in a different context, in that the degree of environmental inducibility of methylation state is (inversely) indicative of the degree of intrinsic regulation. We therefore use the term ‘inducibility’ loosely to refer to the sensitivity of a site to methylation change in response to the environment, regardless of its potential adaptive importance. We found that elevated π of DMCs in general, as well as the elevated Fst of FW-hypo sites, was driven by sites that were environmentally inducible, further supporting a hypothesis of relaxed regulation and relaxed selective constraint at sites that are responsive to environmental conditions. Furthermore, the gain in π among FW-hypo sites was associated with increased environmental inducibility of these sites, suggesting that nucleotide diversity is more likely to accumulate at sites where intrinsic control of methylation is relaxed (and therefore more sensitive to the environment). Indeed, the positive correlation between π and the degree of inducibility suggests that the more sensitive the methylation state is to the environment, the more likely mutations are to be selectively neutral. Therefore, shifts in inducibility (in addition to shifts in methylation variance, as discussed above) may precede shifts in nucleotide diversity. Our results suggest that the majority of environmentally inducible sites are simply ‘blowing in the wind’ and do not have important functions for plasticity which would constrain nucleotide diversity. Nevertheless, in their analysis of Baltic Sea sticklebacks, Heckwolf et al. (2020) observed that the Fst of induced sites (marine fish responsive to lower salinity) depended on the direction of the induced change. Sites that were induced to the ‘evolved’ methylation state observed in the derived freshwater population had lower Fst than those that were induced in the opposite direction. This suggests that some environmentally inducible sites are indeed constrained by selection due to the importance of site plasticity. Here, we did not consider the direction of inducible change, merely considering inducibility as a proxy for the relative weakness of intrinsic regulation.

Again, these observations could also be reconciled with the scenario of a soft sweep, as it is also possible that plasticity of only some methylation sites is necessary to confer adaptation. In other words, plasticity of multiple sites provides multiple alternate routes to adaptation. As many methylation sites would therefore be redundant, they could be lost to mutation without detrimentally affecting the organism’s capacity for adaptive plasticity.

### Limitations and Future directions

Our analyses have revealed striking associations between genetic and epigenetic variation in divergent stickleback populations. However, they focus on one population pair. It is therefore currently not known whether such patterns occur more broadly across different local adaptations (in stickleback and other species) or whether they are idiosyncratic to the relatively recent colonisation event considered in this study (∼700 years). The existence of far older populations, such as those in the Japanese archipelago which are estimated to have colonised freshwater ∼170,000 years ago (Kakioka et al. 2020), raises the question as to the fate of differential methylation over longer periods. Over time, for example, the initially heightened methylation variance may return to a less variable state due to refinement of methylation states via selection or the removal of the CpG sites via accumulation of C-T transitions. Alternatively, no substantial accumulation of mutations over time would suggest that the heightened diversity of FW-hypo sites reflects SGV.

If, indeed, heightened methylation variance arises due to relaxed control of methylation state, the mechanisms by which this could occur are not known. Artemov et al (2017) suggested that mutations in genes encoding epigenetic regulators may underlie increased methylation variance, but did not identify any known epigenetic regulators in the vicinity of genomic regions differentiating marine and freshwater populations in the White Sea region. *Trans*- and *cis*-meQTL have however been identified in stickleback (Hu et al. 2021), some of which are indeed in the vicinity of genomic regions of high Fst between marine and freshwater populations. Differential selection on *trans*-meQTL in particular could have knock on effects on methylation sites across the genome.

Another limitation is that the RRBS data we used came from only a single tissue type (gill), and therefore we were unable to determine which methylation differences between populations are tissue-specific and which are organism-wide. However, many divergent methylation changes are not tissue specific, as a recent analysis of divergent cichlid ecotypes showed that a high proportion of DMRs were shared across tissue types (Vernaz et al. 2021). Also, with respect to the studied divergence between marine and freshwater environments, gills are key to salt homeostasis and their ability to respond to changes osmolarity affects the entire organism.

Finally, the detection of differential methylation is highly sensitive to the analytical methods applied. Namely, the retention of individual-specific CpG sites can lead to the detection of differential methylation simply due to differences in the abundance of CpGs available to be methylated – i.e. directly due to SNPs (Wulfridge et al. 2019). We suggest that whether or not individual-specific CpGs are retained in an analysis, and by extension the definition of ‘differential methylation’, should depend on the goals of the study. Here, by excluding individual-specific sites we could be confident that the detected differential methylation arose through the differential action of the methylation machinery and not due directly to nucleotide variation at the sites themselves. The finding that these sites are nevertheless enriched for SNPs in the population is therefore all the more striking. We acknowledge that excluding individual-specific CpGs risks ignoring a potentially high proportion of methylation variation and, while it was not the goal of this study to extensively characterise this variation, the results should be interpreted with this in mind.

### Conclusions

By intersecting genetic and epigenetic data from naturally diverging populations, we have identified signatures of differential selection on DNAme sites which, combined with patterns of methylation variance and environmental inducibility, support a hypothesis that differential methylation is driven by shifts in the degree of intrinsic control of methylation state in a derived population. Shifts in this control seem to precede increases in nucleotide diversity and may therefore indicate parts of the genome where diversification is imminent. Heightened diversity of DMCs may also reflect a soft sweep which retains diversity at both genetic and epigenetic levels, a scenario compatible with previous genetic studies of local adaptation in stickleback (Terekhanova et al. 2014; Roberts Kingman et al. 2021). Indeed, our results support the idea that epigenetic variation should be incorporated alongside genetic mechanisms of adaptation in models of species adaptation and evolutionary potential (Bernatchez 2016). Our analyses further demonstrate the exciting potential held in published datasets for exploring combined patterns of genome and epigenome evolution.

## Methods

### Datasets

We obtained a reduced-representation bisulfite sequencing (RRBS) dataset published by Artemov et al. (2017) (SRA project accession: PRJNA324599) comprising a total of six marine sticklebacks (of which three were exposed to lower salinity) and six freshwater sticklebacks (of which three were exposed to higher salinity; however, one of these three samples was excluded due to incomplete bisulfite conversion). The freshwater fish used for RRBS were sampled from Lake Mashinnoye, while the marine fish were sampled from the Kandalaksha gulf. Freshwater fish were also collected for pool-seq by Terekhanova et al. (2014) from Lake Mashinnoye (SRA run accession: SRR869609), while marine fish were collected from the Kandalaksha gulf as part of the 2014 study and a subsequent 2019 study (Terekhanova et al. 2019). We selected the ‘White Sea, WSBS’ sample from Terekhanova et al. (2019) (SRR7470095) as the marine sample for our comparison, given that it has a similar pool size to the Mashinnoye sample (12 vs 10) and a similar number of reads (64,176,648 vs 62,016,859 after quality trimming). Sequence files were obtained in FASTQ format from the Sequence Read Archive (SRA) and European Nucleotide Archive (ENA).

### Data processing: RRBS

Raw RRBS reads were trimmed using TrimGalore v0.6.6 using default settings. Alignment to the Threespine stickleback v.5 assembly (Nath et al. 2021) and subsequent methylation calling were carried out using Bismark v0.22.3 (Krueger and Andrews 2011) with Bowtie2 v2.3.4.1 as the aligner (Langmead and Salzberg 2012). Methylation calls were not strand-specific. To remove sites harbouring C-T/G-A SNPs which otherwise contribute erroneous counts of non-methylated Cs, we ran BS-SNPer v1.1 (Gao et al. 2015) on each sample using a minimum coverage of 5 and otherwise default settings, compiled the coordinates of all sites harbouring C-T/G-A SNPs in any of the individuals (either homo- or heterozygous), and removed these sites from the Bismark coverage files containing the methylation counts (counts of Cs and Ts at each position).

### Data processing: Pool-seq

Raw reads were trimmed with Trimmomatic v0.36 (Bolger et al. 2014) with the option SLIDINGWINDOW:4:20 and otherwise default parameters. Only reads which remained paired after trimming were kept. Reads were mapped to the Threespine stickleback v.5 assembly with Bowtie2 v2.3.4.1 with default parameters. Sambamba v0.7.1 (Tarasov et al. 2015) was used to filter the alignments to retain those with MAPQ >/=20 and to remove PCR duplicates. Samtools v0.1.18 (Danecek et al. 2021) was used to generate a pileup file from each BAM file, as required for the Popoolation and Popoolation2 toolkits.

### Identification of differentially methylated CpG sites (DMCs) and subsampling of non-DMCs

Site-level differential methylation analysis was carried out using the methylKit R package v1.22.0 (Akalin et al. 2012), inputting the SNP-filtered Bismark coverage files. We omitted sites which did not have at least 5x coverage in each of the 11 samples as well as sites located on the mitochondrial chromosome and two sex chromosomes (chromosomes XIX and Y). We then filtered out sites that had either 0% or 100% methylation (i.e. no variation) in all samples from the main population comparison (3x marine and 3x freshwater). Three differential methylation analyses were then performed separately, comprising the comparisons also described in Artemov et al (2017): (1) marine fish in saltwater vs freshwater fish in freshwater (main population comparison), (2) marine fish in saltwater vs marine fish in freshwater (marine low salinity treatment), and (3) freshwater fish in freshwater vs freshwater fish in saltwater (freshwater high salinity treatment). All groups comprised N=3 with the exception of freshwater fish in saltwater (N=2, due to low bisulfite conversion efficiency of sample SRR3632642). Regardless, we considered sites to be differentially methylated given a change in % methylation of >/= 15 and a FDR-corrected p-value of </= 0.05. The purpose of the experimental comparisons was to identify which population-DMCs were also induced by salinity change, and so we did not consider sites that were induced by salinity change but not differentially methylated between populations. We also did not consider the direction of induced change (hypo- or hypermethylated in response to salinity change). Subsequently, we detected 92,710 sites that were hypomethylated in freshwater compared to marine (of which 10,765 were induced in either population) and 15,057 sites that were hypermethylated in freshwater compared to marine (of which 6063 were induced in either population). 897,733 sites were not differentially methylated between populations, which we would use as reference sites when examining nucleotide diversity. Due to the possibility that DMCs could be distributed non-randomly across a chromosome and given that nucleotide diversity can vary across a chromosome (e.g. lower diversity in centromeric regions), we used a sampling procedure which randomly selected one non-differentially methylated site that was within 2kb upstream or 2kb downstream of each DMC. After removing duplicate samples, this resulted in a subsample of 100,887 non-differentially methylated CpG sites (non-DMCs).

### Nucleotide diversity of site categories

We used the variance-at-position.pl script from the Popoolation toolkit (Kofler, Orozco-terWengel, et al. 2011) to calculate within-population nucleotide diversity statistics (π, Watterson’s *θ*, and Tajima’s *D*) for different categories of site (e.g. non-DMC, FW-hypo, FW-hyper) on each chromosome for each population. The decision to obtain estimates of π on a per-chromosome basis was because Popoolation’s estimates of π are accurate over large numbers of sites, but not at the single site level (Kofler, Orozco-terWengel, et al. 2011). Each site was labelled with its chromosome and its category within the analysis (e.g. chrI FW-hypo) and the labelled category was entered as the ‘gene ID’ in a GTF file, such that variance-at-position.pl, which was developed to calculate diversity statistics per-gene, was instructed to calculate π for each combination of chromosome and site category. A similar procedure was used to obtain π for sites ranked according to (change in) mean % methylation, (change in) SD of % methylation, and absolute inducibility, whereby ranks were assigned using the bin() function from the OneR package and sites were labelled in the GTF according to their rank (regardless of chromosome), such that a single value of π was obtained for each rank. Variance-at-position.pl from Popoolation was run with the parameters --min-qual 20 --min-coverage 3 --min-count 2.

Fst for different categories of methylation sites (including ranked sites) were obtained using the Popoolation2 toolkit (Kofler, Pandey, et al. 2011). ‘Gene-wise’ .sync files were obtained from the pileup files using coordinates in the abovementioned gtf files and were used as input for the ‘fst-sliding.pl’ script which was run with parameters --min-count 2 --min-coverage 3 -- pool-size 22 --min-covered-fraction 0 --max-coverage 1000 --window-size 1000000 --step-size 1000000.

### Percentage of sites with SNPs

To obtain the % of sites within each result category (non-DMC, FW-hypo, and FW-hyper) harbouring SNPs of different types (C-T/G-A and non-C-T/G-A) in the pool-seq samples, we filtered the BAM files of each population to retain alignments corresponding with the positions of interest. We then ran GATK HaplotypeCaller (McKenna et al. 2010) with the –sample-ploidy set to the pool size x 2 (24 for marine and 20 for freshwater), and otherwise default settings. The subsequent VCF file was then filtered using bcftools v1.10 to retain only biallelic SNPs at the sites of interest. We subsequently extracted from the VCF a list of reference and alternate alleles at sites of interest harbouring biallelic SNPs. We were therefore able to assign SNPs as either ‘C-T/G-A’ or ‘other’, and calculate the % of sites in each result category harbouring biallelic SNPs of one of those two classes.

### Statistical comparisons

All plotting and statistical analyses were carried out in R version 4.2.0 (R Development Core Team 2011), with plots generated using the ggplot2 package (Wickham 2011). For comparing nucleotide diversity between populations, we first used Shapiro-Wilkes tests to determine whether the distribution of pairwise differences (paired chromosomes) differed significantly from a normal distribution. Paired *T*-tests were used in the case of normally-distributed pairwise differences and paired Wilcoxon tests were used in the case of non-normally-distributed pairwise differences. Linear models of nucleotide diversity parameters as a function of inducibility were fit using the lm() function from the stats package.

## Supporting information

BioRender license

BioRender license

## Abbreviations

DNAme: DNA methylation
DM: Differential Methylation
DMC: Differentially Methylated Cytosine
DMR: Differentially Methylated Region
FW: Freshwater
FW-hyper: Hypermethylated in Freshwater population
FW-hypo: Hypomethylated in Freshwater population
Non-DMC: Non-Differentially Methylated Cytosine
RRBS: Reduced Representation Bisulfite Sequencing
SD: Standard Deviation
SGV: Standing Genetic Variation
SNP: Single Nucleotide Polymorphism

## Supplementary material

Supplementary figures and methods are available in the online supplementary material.

## Acknowledgements

The authors are grateful to Katie Peichel and Dragan Stajic for insightful discussion. Calculations were performed on UBELIX (http://www.id.unibe.ch/hpc), the HPC cluster at the University of Bern. Figures 1 and 5 were created with BioRender.com. This project has received funding from the European Research Council (ERC) under the European Union’s Horizon 2020 research and innovation programme Grant agreement No. 947636.

## Data availability

This work used entirely pre-existing datasets available under SRA project accessions PRJNA324599 (RRBS data), PRJNA204958, and PRJNA479509 (whole genome pool-seq). Code for performing analyses has been deposited on GitHub in the repository: https://github.com/jamesord/StickMethDiv.

## Notes

https://github.com/jamesord/StickMethDiv

https://www.ncbi.nlm.nih.gov/bioproject/PRJNA324599/

https://www.ncbi.nlm.nih.gov/bioproject/PRJNA204958

https://www.ncbi.nlm.nih.gov/bioproject/PRJNA479509

### Competing Interest Statement

The authors have declared no competing interest.

### Summary of Updates

No alterations to intro, methods, or results. Some changes to the discussion (improved clarity and slight restructuring), but conclusions are unchanged. Some small changes to the abstract.

