## Supplementary material for "High nucleotide diversity accompanies differential DNA methylation in naturally diverging populations"

### Supplementary figures

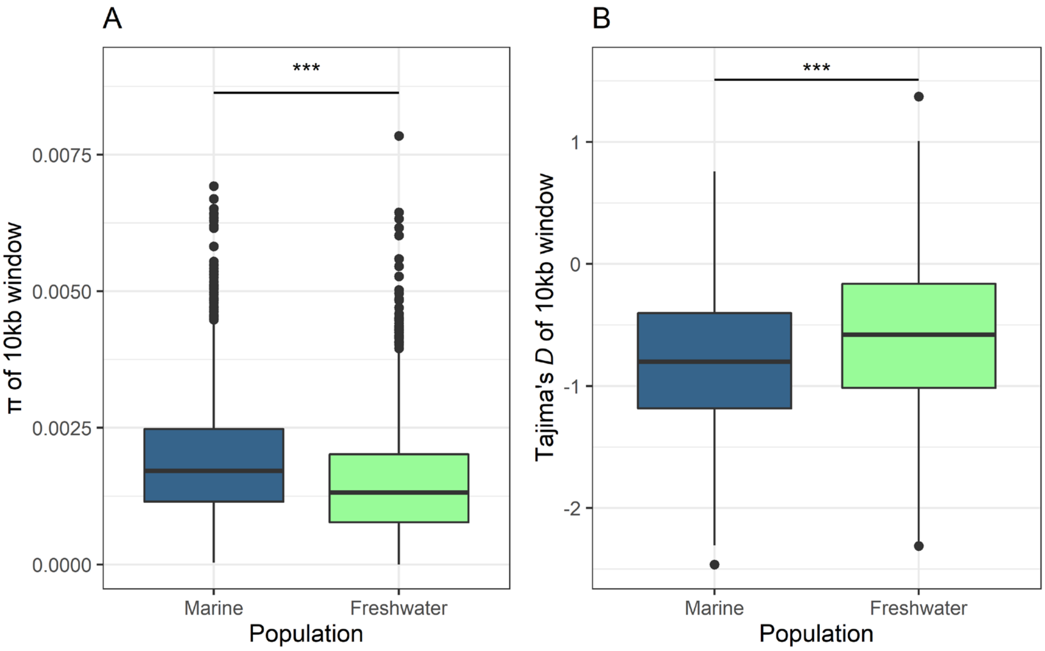


**Figure S1. Nucleotide diversity statistics of sliding windows across chromosome I of marine and freshwater population.** Estimates of π and Tajima's *D* in 10kb windows across chromosome I were derived from pool-seq data of marine (blue) and freshwater (green) populations. Significance stars represent results of general independence tests with approximated (Monte Carlo) null distributions and one-sided hypotheses.

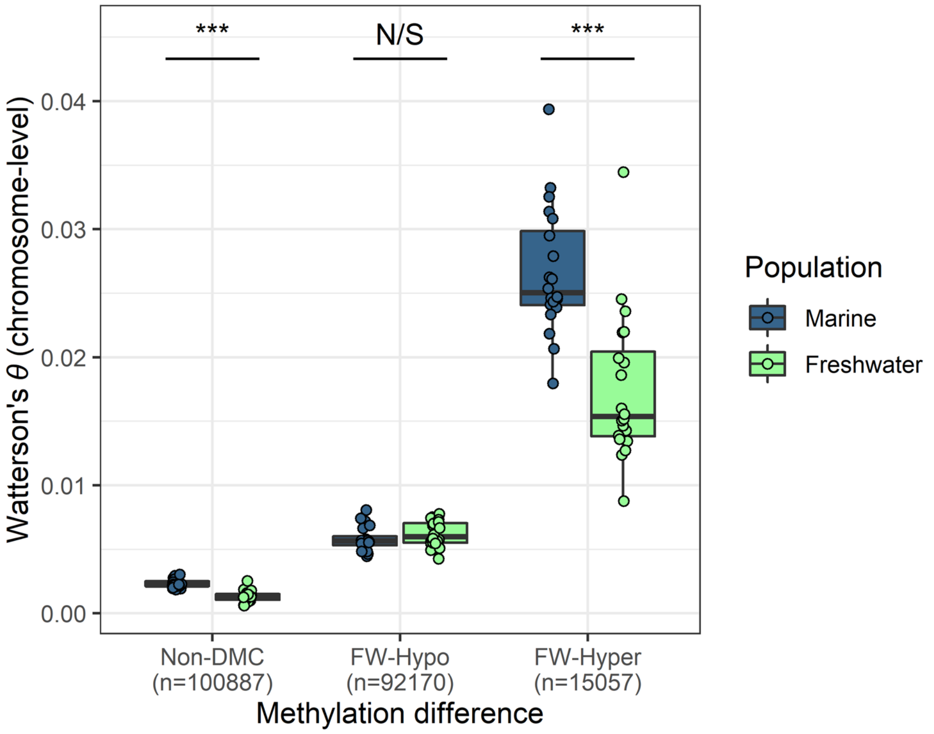


**Figure S2. Watterson's $\theta$ estimated from pool-seq of marine and freshwater sticklebacks for three classes of methylation site.** Sites were classified according to the direction of methylation difference in freshwater fish compared to marine. $\theta$ was estimated for each chromosome separately, such that one point represents an estimate from one chromosome. Significance stars derive from paired Wilcoxon tests (comparison of chromosome pairs).

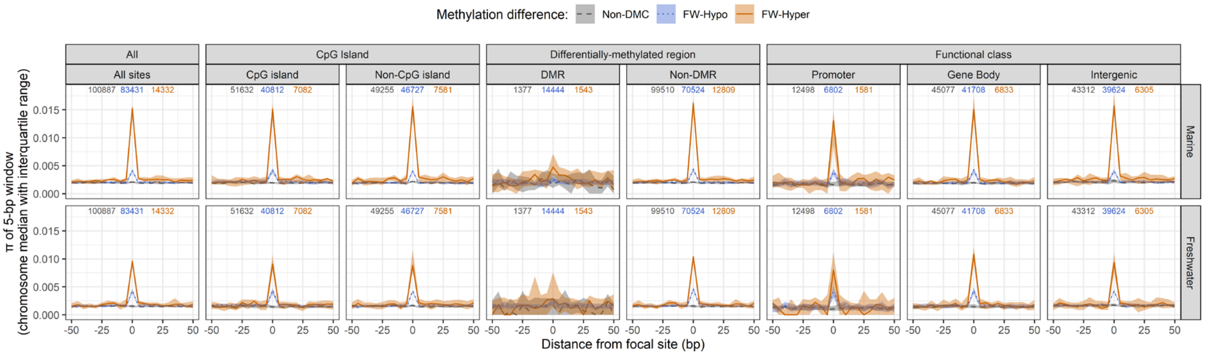


**Figure S3. Nucleotide diversity surrounding different classes of methylation site.** π was calculated in windows of 5bp with steps of 5bp extending 50bp either side of the focal site. Sites are classified according to the direction of methylation difference in freshwater fish compared to marine (non-DMC, dashed black lines; FW-hypo, blue dotted lines; FW-hyper, solid orange lines). Subplots are divided by population (marine, top and freshwater, bottom) and the category of genomic feature in which sites are considered (from left to right: all sites, sites within CpG islands, not within CpG islands, within differentially methylated regions (DMRs), not within DMRs, within promoter regions, within gene bodies, and within intergenic regions). Separate sets of sliding windows were derived for each chromosome and therefore the middle lines and ribbons denote the median and interquartile range of all chromosome-level estimates, respectively. The total numbers of non-DMC, FW-hypo sites, and FW-hyper sites considered in each category (total across all chromosomes) are shown at the top of each panel in grey, blue, and orange, respectively.

### Supplementary methods

*Sliding windows along chromosome I*

To obtain estimates of π and Tajima's *D* for each population along chromosome I, the pileup files generated from the WSBS and Mashinnoye pool-seq alignments were subset to include only data from chromosome I alignments. The variance-sliding.pl script from Popoolation (Kofler et al. 2011) was then run on each of the aforementioned pileup files. Parameters used for π estimation were: --window-size 10000 --step-size 10000 --min-count 2 --min-coverage 4 --min-qual 20. Parameters used for Tajima's *D* estimation were: -window-size 10000 --step-size 10000 --min-count 1 --min-coverage 3 --min-qual 20 --dissable-corrections on.

*Sliding windows around sites within different genomic features*

To visualise π in windows around DMCs in different genomic features, we first obtained the coordinates of genes (from the gff3 annotation file provided at https://stickleback.genetics.uga.edu), promoters (classified arbitrarily as the region spanning from 1kb upstream to 0.5kb downstream of the gene start; Heckwolf et al., 2020), CpG islands, and differentially methylated regions (DMRs). CpG islands were identified from the stickleback v.5 assembly (Nath et al. 2021) by running CpGIscan (https://github.com/jzuoyi/cpgiscan) with default parameters. DMRs (differentially methylated regions between marine and freshwater fish) were identified from the SNP-filtered Bismark coverage using the R package bsseq v1.32.0 (Hansen et al. 2012). Loci were retained for DMR calling if they had at least 5x coverage in each of the six samples (3x marine and 3x freshwater), and a region was considered differentially methylated given a T-stat of >/= 1.5 and a mean difference in % methylation of >/= 15. Subsequently, the coordinates of all aforementioned genomic features were compiled as BED files, such that for each feature type, a BED file containing the sites of interest (DMCs plus the subsample of non-DMCs) could be filtered using bedtools-intersect from bedtools v2.29.2 (Quinlan and Hall 2010) to return a list of the sites that were within that feature. We subsequently obtained lists of sites assigned to eight feature categories: 'all sites' (regardless of feature), 'CpG islands', 'non-CpG islands', 'DMRs', 'non-DMRs', 'promoters', 'genes', and 'intergenic'. We considered 'intergenic' sites to be any sites that did not overlap with either promoters or genes. For each site within each feature, we established a 5bp window centred on the site which was labelled as 0, and ten 5bp windows in steps of 5bp each side of the site labelled from -50 to 50. Window labels were then entered into GTF files for use with variance-at-position.pl from Popoolation (Kofler et al. 2011), such that a single π value was calculated for each window, encompassing all of the sites in a given category. The above was carried out separately for each combination of result category (non-DMC, FW-hypo, and FW-hyper), genomic feature category, and chromosome.
